# Broad Host Range of SARS-CoV-2 Predicted by Comparative and Structural Analysis of ACE2 in Vertebrates

**DOI:** 10.1101/2020.04.16.045302

**Authors:** Joana Damas, Graham M. Hughes, Kathleen C. Keough, Corrie A. Painter, Nicole S. Persky, Marco Corbo, Michael Hiller, Klaus-Peter Koepfli, Andreas R. Pfenning, Huabin Zhao, Diane P. Genereux, Ross Swofford, Katherine S. Pollard, Oliver A. Ryder, Martin T. Nweeia, Kerstin Lindblad-Toh, Emma C. Teeling, Elinor K. Karlsson, Harris A. Lewin

## Abstract

The novel coronavirus SARS-CoV-2 is the cause of Coronavirus Disease-2019 (COVID-19). The main receptor of SARS-CoV-2, angiotensin I converting enzyme 2 (ACE2), is now undergoing extensive scrutiny to understand the routes of transmission and sensitivity in different species. Here, we utilized a unique dataset of 410 vertebrates, including 252 mammals, to study cross-species conservation of ACE2 and its likelihood to function as a SARS-CoV-2 receptor. We designed a five-category ranking score based on the conservation properties of 25 amino acids important for the binding between receptor and virus, classifying all species from *very high* to *very low*. Only mammals fell into the *medium* to *very high* categories, and only catarrhine primates in the *very high* category, suggesting that they are at high risk for SARS-CoV-2 infection. We employed a protein structural analysis to qualitatively assess whether amino acid changes at variable residues would be likely to disrupt ACE2/SARS-CoV-2 binding, and found the number of predicted unfavorable changes significantly correlated with the binding score. Extending this analysis to human population data, we found only rare (<0.1%) variants in 10/25 binding sites. In addition, we observed evidence of positive selection in ACE2 in multiple species, including bats. Utilized appropriately, our results may lead to the identification of intermediate host species for SARS-CoV-2, justify the selection of animal models of COVID-19, and assist the conservation of animals both in native habitats and in human care.

## Introduction

The 2019-novel coronavirus (2019-nCoV, also, SARS-CoV-2 and COVID-19 virus) is the cause of Coronavirus Disease-2019 (COVID-19), a major pandemic that threatens millions of lives and the global economy (1). Comparative analysis of SARS-CoV-2 and related coronavirus sequences has shown that SARS-CoV and SARS-CoV-2 likely originated in bats, followed by transmission to an intermediate host, and that both viruses may have an extended host range that includes primates and other mammals (1–3). However, the immediate source population/species for SARS-CoV and SARS-CoV-2 viruses has not yet been identified. Several mammalian species host coronaviruses, and these infections are frequently associated with severe clinical diseases, such as respiratory and enteric disease in pigs and cattle (4, 5). Molecular phylogenetics revealed that at least one human coronavirus (HCov-OC43), may have originated in cattle or swine (6), and that this virus was associated with a human pandemic that emerged in the late 19^th^ century (7). Recent data indicate that coronaviruses can move from bats to other wildlife species and humans (8) and from humans to tigers (9) and pigs (10). Therefore, understanding the host range of SARS-CoV-2 and related coronaviruses is essential for improving our ability to predict and control future pandemics. It is also crucial for protecting populations of wildlife species in native habitats and under human care, particularly non-human primates, who may also be susceptible to COVID-19 (11).

The angiotensin I converting enzyme 2 (ACE2) serves as a functional receptor for the spike protein (S) of SARS-CoV and SARS-CoV-2 (12, 13). Under normal physiological conditions, ACE2 is a dipeptidyl carboxypeptidase that catalyzes the conversion of angiotensin I into angiotensin 1-9, a peptide of unknown function, and angiotensin II, a vasoconstrictor that is important in the regulation of blood pressure (14). ACE2 also converts angiotensin II into angiotensin 1-7, a vasodilator that affects the cardiovascular system (14) and may regulate other components of the renin-angiotensin system (15). The host range of SARS-CoV-2 may be extremely broad due to the conservation of ACE2 in mammals (2, 13) and its expression on ciliated bronchial epithelial cells and type II pneumocytes (10). While coronaviruses related to SARS-CoV-2 use ACE2 as a primary receptor, coronaviruses may use other proteases as receptors, such as CD26 (DPP4) for MERS-CoV (16), thus limiting or extending their host range.

In humans, ACE2 may be a cell membrane protein or it may be secreted (14). The secreted form is created primarily by enzymatic cleavage of surface-bound ACE2 by ADAM17 and other proteases (14). Sequence variation in *ACE2* affects the protein’s functions. *ACE2* is polymorphic in humans, with many synonymous and nonsynonymous mutations identified, although most are rare at the population level (17) and few are believed to affect cellular susceptibility to human coronavirus infections (18). Site-directed mutagenesis and co-precipitation of SARS-CoV constructs have revealed critical residues on the ACE2 tertiary structure that are essential for binding to the virus receptor binding domain (RBD) (19). These findings have been strongly supported by co-crystallization and the structural determination of the SARS-CoV and SARS-CoV-2 S proteins with human ACE2 (13, 20, 21), as well as binding-affinity with heterologous ACE2 (19). The RBD of human coronaviruses may mutate to change the binding affinity of S for ACE2, and thus lead to adaptation in humans or other hosts. The best studied example is the palm civet, believed to have been the intermediate host between bats and humans for SARS-CoV (2). To date, an intermediate host for SARS-CoV-2 has not been identified definitively, although Malayan pangolins (*Manis javanica*) have been proposed as a possible reservoir (22).

Comparative analysis of ACE2 nucleotide and protein sequences can predict their ability to bind SARS-CoV-2 S and therefore will yield important insights into the biology and potential zoonotic transmission of SARS-CoV-2 infection. Recent work has examined ACE2 from different vertebrate species and predicted its ability to bind SARS-CoV-2 S, but phylogenetic sampling was extremely limited (11, 23). Here, we made use of sequenced genomes of 410 vertebrates and protein structural analysis, to identify ACE2 homologs in all vertebrate classes (fishes, amphibians, birds, reptiles, and mammals) that have the potential to serve as a receptor for SARS-CoV-2, and to understand the evolution of ACE2 SARS-CoV-2 S binding sites. Our results reinforce earlier findings on the natural host range of SARS-CoV-2, and predict a broader group of species that may serve as a reservoir or intermediate host for this virus. Importantly, many threatened and endangered species were found to be at potential risk for SARS-CoV-2 infection, suggesting that as the pandemic spreads, humans could inadvertently introduce a potentially devastating new threat to these already vulnerable populations, especially for great apes and other primates.

## Results

### Comparison of vertebrate ACE2 sequences and their predicted ability to bind SARS-CoV-2

We identified 410 unique vertebrate species with *ACE2* orthologs (Dataset S1) that included representatives of all vertebrate taxonomic classes. Among these were 252 mammals, 72 birds, 65 fishes, 17 reptiles and 4 amphibians. Twenty-five amino acids corresponding to known SARS-CoV-2 S-binding residues (11, 13, 21) were examined for their similarity to the residues in human ACE2 (Fig. 1, Dataset S1). On the basis of known interactions between specific residues on ACE2 and the RBD of SARS-CoV-2 S, a set of rules was developed for predicting the likelihood of S binding to ACE2 from each species (see Materials and Methods). Five score categories were predicted: *very high*, *high*, *medium*, *low* and *very low*. Results for all species and all SARS-CoV-2 S binding scores are shown in Dataset S1, and results for mammalian species are also shown in Fig. 1. The *very high* classification had at least 23/25 ACE2 residues identical to their human homolog and other constraints on substitutions at SARS-CoV-2 S binding hot spots (see Materials and Methods). The 18 species predicted as *very high* were all Old World primates and apes completely identical to human across the 25 ACE2 binding residues. The ACE2 proteins of 28 species were classified as having a *high* likelihood of binding the S RBD. Among them are twelve cetaceans, seven rodents, three cervids (deer), three lemuriform primates, two representatives of the order Pilosa (Giant anteater and Southern tamandua), and one Old World primate (Angola colobus, Fig. 1). Fifty-seven species scored as *medium* for the ability of their ACE2 to bind SARS-CoV-2 S. Like the *high* score, this category has at least 20/25 residues identical to human ACE2 but more relaxed constraints for critical binding residues. All species with *medium* score are mammals distributed across six orders.

**Figure 1.**
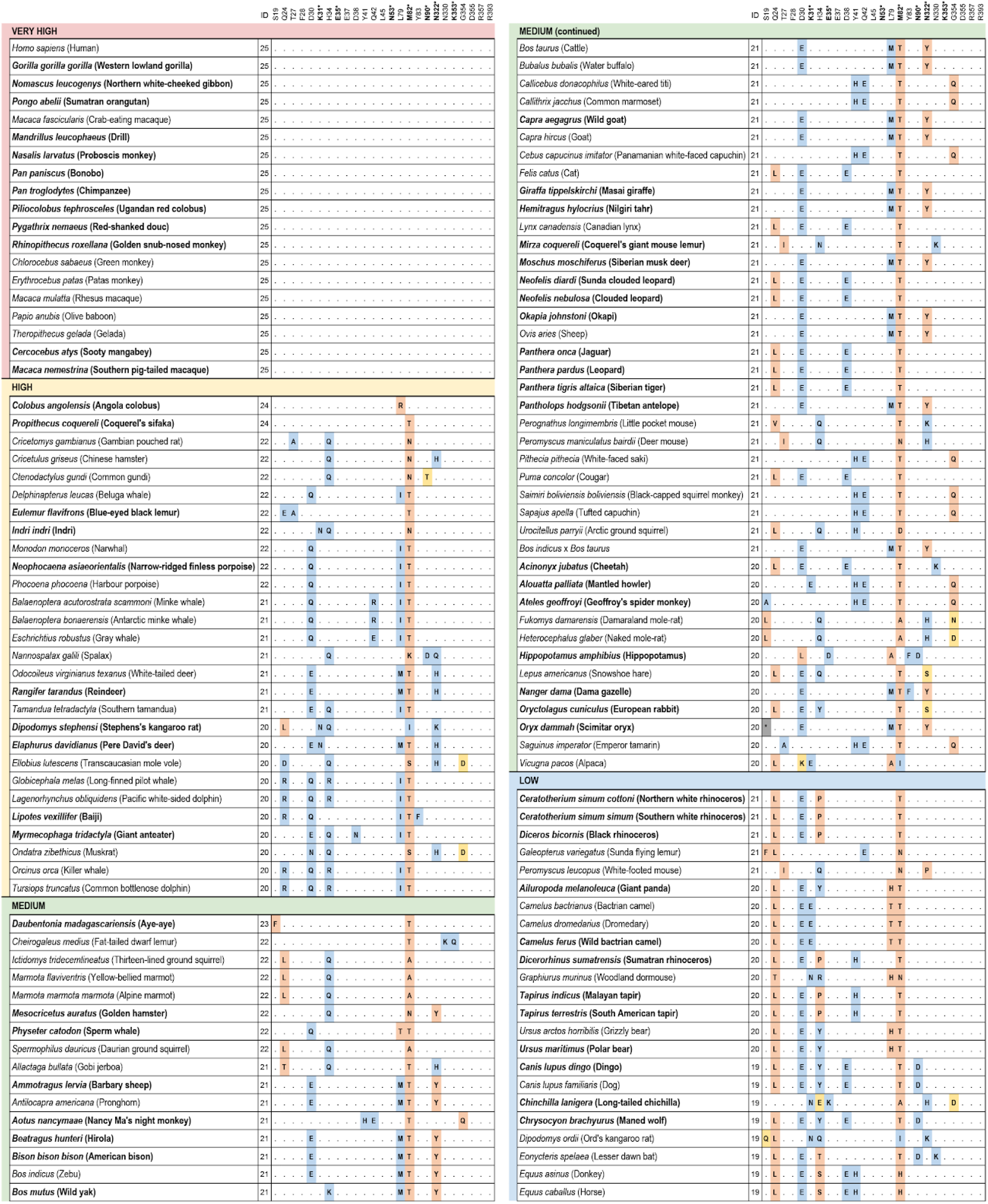
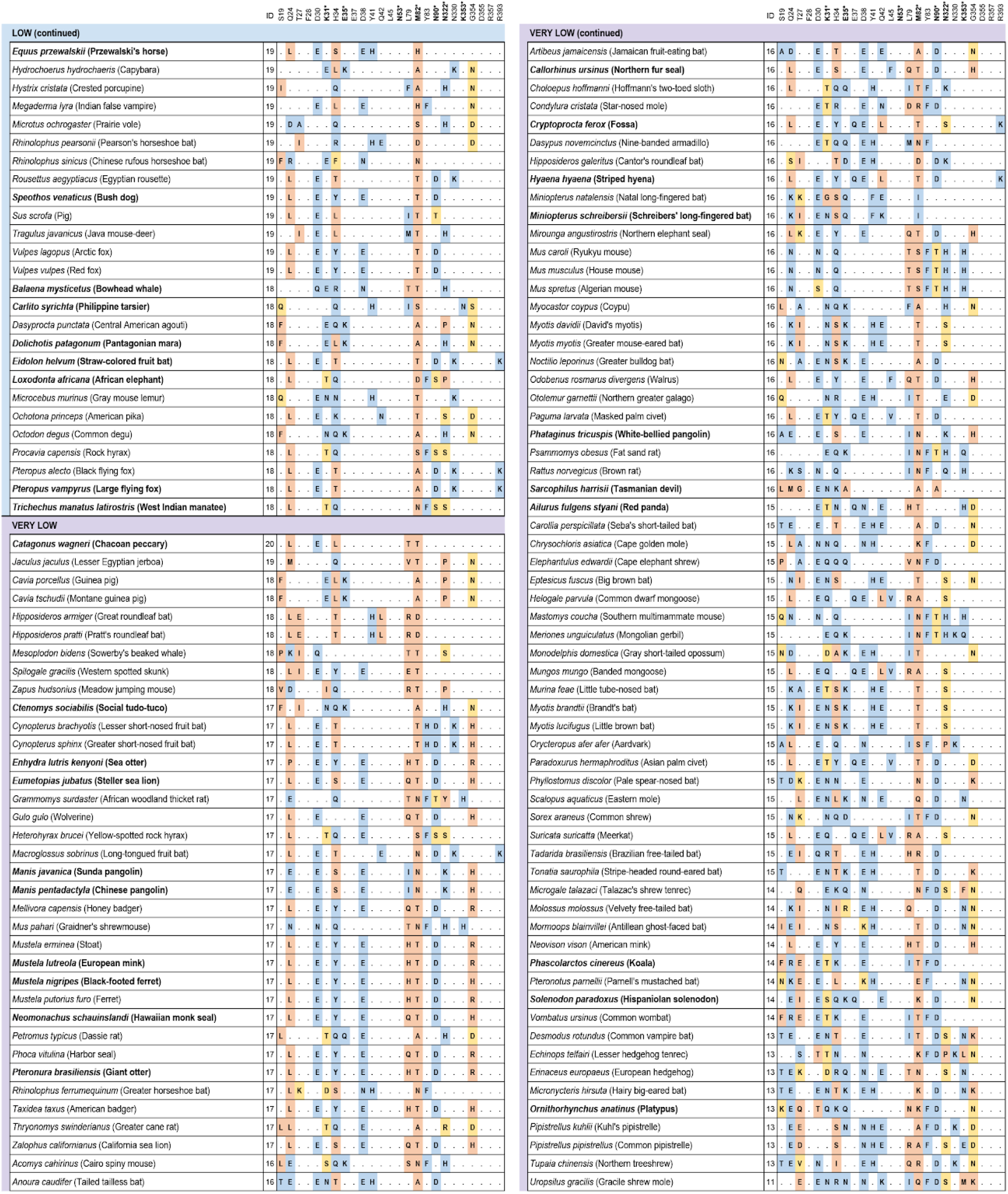
Cross-species conservation of ACE2 and predictions of SARS-CoV-2 susceptibility. Species are sorted by binding score of ACE2 for SARS-CoV-2 S. The ‘ID’ column depicts the number of amino acids identical to human binding residues. Bold amino acid positions (also labeled with *) represent residues at binding hotspots and constrained in the scoring scheme. Each amino acid substitution is colored according to its classification as non-conservative (orange), semi-conservative (yellow) or neutral (blue), as compared to the human residue. Bold species names depict species with threatened IUCN risk status. The 410 vertebrate species dataset is available in Dataset S1.

Among Carnivora, 9/43 scored *medium*, 9/43 scored *low*, and 25/43 scored *very low* (Fig. 1). The carnivores scoring *medium* were only felids, including the domestic cat and Siberian tiger. Among the 13 Primates scoring *medium* there were 10 New World primates and three lemurs. Of 45 Rodentia species, 11 scored *medium*. Twenty-one Artiodactyls scored *medium*, including several important wild and domesticated ruminants, such as domesticated cattle, bison, sheep, goat, water buffalo, Masai giraffe, and Tibetan antelope. Species scoring *medium* also included 2/3 Lagomorphs and one Cetacean (sperm whale).

All chiropterans (bats) scored *low* (N=8) or *very low* (N=29) (Fig. 1), including the Chinese rufous horseshoe bat (*Rhinolophus sinicus*), from which a coronavirus very similar to SARS-CoV-2 was identified (1). Only 7.7% (3/39) primate species’ ACE2 scored *low* or *very low*, and 61% of rodent species scored *low* (10/46) or *very low* (18/46). All monotremes (N=1) and marsupials (N=4) scored *very low*. All birds, fish, amphibians, and reptiles scored *very low*, with less than 18/25 ACE2 residues identical to the human and many non-conservative residues at the remaining non-identical sites (Dataset S1). Notable species scoring *very low* include the Chinese pangolin (*Manis pentadactyla*), Sunda pangolin (*Manis javanica*), and white-bellied pangolin (*Phataginus tricuspis*) (Fig. 1, Dataset S1).

### Structural analysis of the ACE2/SARS-CoV-2 S binding interface

We complemented the sequence-identity based scoring scheme with a qualitative approach that combined structural homology modeling and best fit rotamer positioning. We examined the 25 ACE2 binding residues in a subset of 28 representative species (Fig. S1) and 17 sites were variable and not glycosylation sites. First, we assessed the similarity of every contact at the binding interface between two recently solved crystal structures for the human ACE2/SARS-CoV-2 S RBD complex in humans, 6M0J and 6WV1 (13, 21). Both structures were in agreement except for the position of S19, which was excluded from subsequent analysis (24). We then generated homology models, and aligned them to the human ACE2/SARS-CoV-2 S RBD 6M0J structure. This showed a high degree of similarity along the C⍺ backbone (25) for each of the 28 species. We selected the most favorable rotamer at each residue using CHIMERA (Fig. S2).

We examined a total of 55 substitutions and assigned each to one of three types: *neutral* (N; likely to maintain similar contacts; 18 substitutions); *weaken* (W; likely to weaken the interaction; 14 substitutions); or *unfavorable* (U; likely to introduce unfavorable interactions; 23 substitutions) (Fig. S1). Our assignments show good agreement with those made in a second study (26) based on experimental data, with 83.4% of the 55 substitutions evaluated concordant between the two approaches (Fig. S1). The structural homology binding assessments support the sequence identity analysis, with the fraction of residues ranked as U, correlating very strongly with the substitution scoring scheme (Spearman correlation rho=0.76; p< 2.2e-16; Fig. 2).

**Figure 2.**
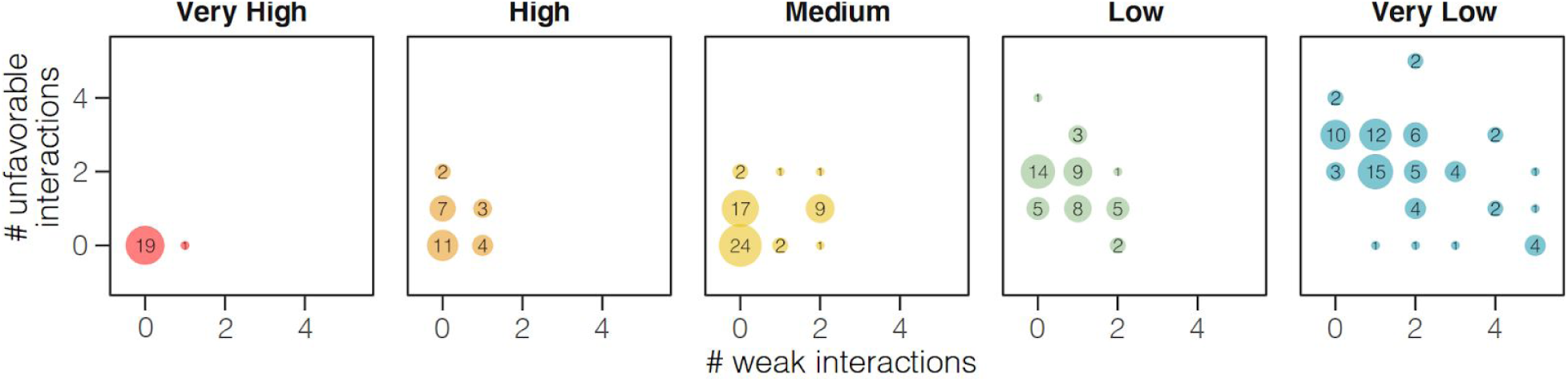
Congruence between binding score and structural homology analysis. Species classified by sequence identity to human ACE2 as *very high* (red) or *high* binding score (orange) have significantly fewer amino acid substitutions rated as potentially altering the binding interface between ACE2 and SARS-CoV-2 through protein structural analysis, as compared to *low* (green) or *very low* (blue) species. The more severe *unfavorable* variants are counted on y-axis and less severe *weaken* variants on the x-axis. Black numerical labels indicate species count.

### Structural analysis of variation in human *ACE2*

We applied the same approach used to compare species, sequence identity and protein structural analysis, to examine the variation in ACE binding residues within humans, some of which have been proposed to alter binding affinity (18, 27–30). We integrated data from six different sources: dbSNP (31), 1KGP (32), Topmed (33), UK10K (34) and CHINAMAP (28), and identified a total of 11 variants in ten of the 25 ACE2 binding residues (Dataset S2). All variants found are rare, with allele frequency less than 0.01 in any populations, and less than 0.0007 over all populations. Three of the 11 variants were synonymous changes, seven were conservative missense variants, and one, S19P, was a semi-conservative substitution. S19P has the highest allele frequency of the 11 variants, with a global frequency of 0.0003 (17). We evaluated, by structural homology, six missense variants. Four were *neutral* and two weakening (E35K, frequency=0.000016; E35D, frequency=0.000279799). S19P was not included in our structural homology assessment, but a recent study predicted it would increase binding affinity (26). Thus, with an estimated summed frequency of 0.001, genetic variation in the ACE2 S-binding interface is overall rare, and it is unclear whether the variation that does exist increases or decreases susceptibility to infection.

### Evolution of ACE2 across mammals

We next investigated the evolution of ACE2 variation in vertebrates, including how patterns of positive selection compare between bats, a mammalian lineage known to harbor a diversity of coronaviruses (35), and other mammalian clades. We first inferred the phylogeny of *ACE2* using our 410-vertebrate alignment and IQTREE, using the best-fit model of sequence evolution (JTT+F+R7) and rooting the topology on fishes (Dataset S3; Fig. S3). We then assayed sequence conservation with PhyloP (36). The majority of ACE2 codons are significantly conserved across vertebrates and across mammals, likely reflecting its critical function in the renin-angiotensin system (37) (Dataset S4.1), with ten residues in the ACE2 binding domain exceptionally conserved in Chiroptera and/or Rodentia (Dataset S4.2).

We next used phyloP and CODEML to test for acceleration and positive selection (36). PhyloP compares the rate of evolution at each codon to the expected rate in a model estimated from third nucleotide positions of the codon, and is agnostic to synonymous versus nonsynonymous substitutions (dN/dS). CODEML uses ⍵=dN/dS>1 and Bayes Empirical Bayes (BEB) scores to identify codons under positive selection, and was run on a subset of 64 representative mammals (see Materials and Methods).

*ACE2* shows significant evidence of positive selection across mammals (⍵=1.83, LRT=194.13, p<0.001; Dataset S4.3, 4.4). Almost 10% of codons (N=73; 9 near the RBD) are accelerated within mammals (Dataset S4.1, 4.5), and 18 of these have BEB scores greater than 0.95, indicating positively selected residues (Dataset S4.5, 4.6, Fig. S4). Nineteen accelerated residues, including two positively-selected codons (Q24, H34), are critical for the binding of the ACE2 RBD and SARS-CoV-2 S (Dataset S4.5; Fig. 3; Fig. S5). Q24 has not been observed to be polymorphic within the human population, and H34 harbors a synonymous polymorphism (AF=0.00063) but no non-synonymous polymorphisms (Dataset S2).

**Figure 3.**
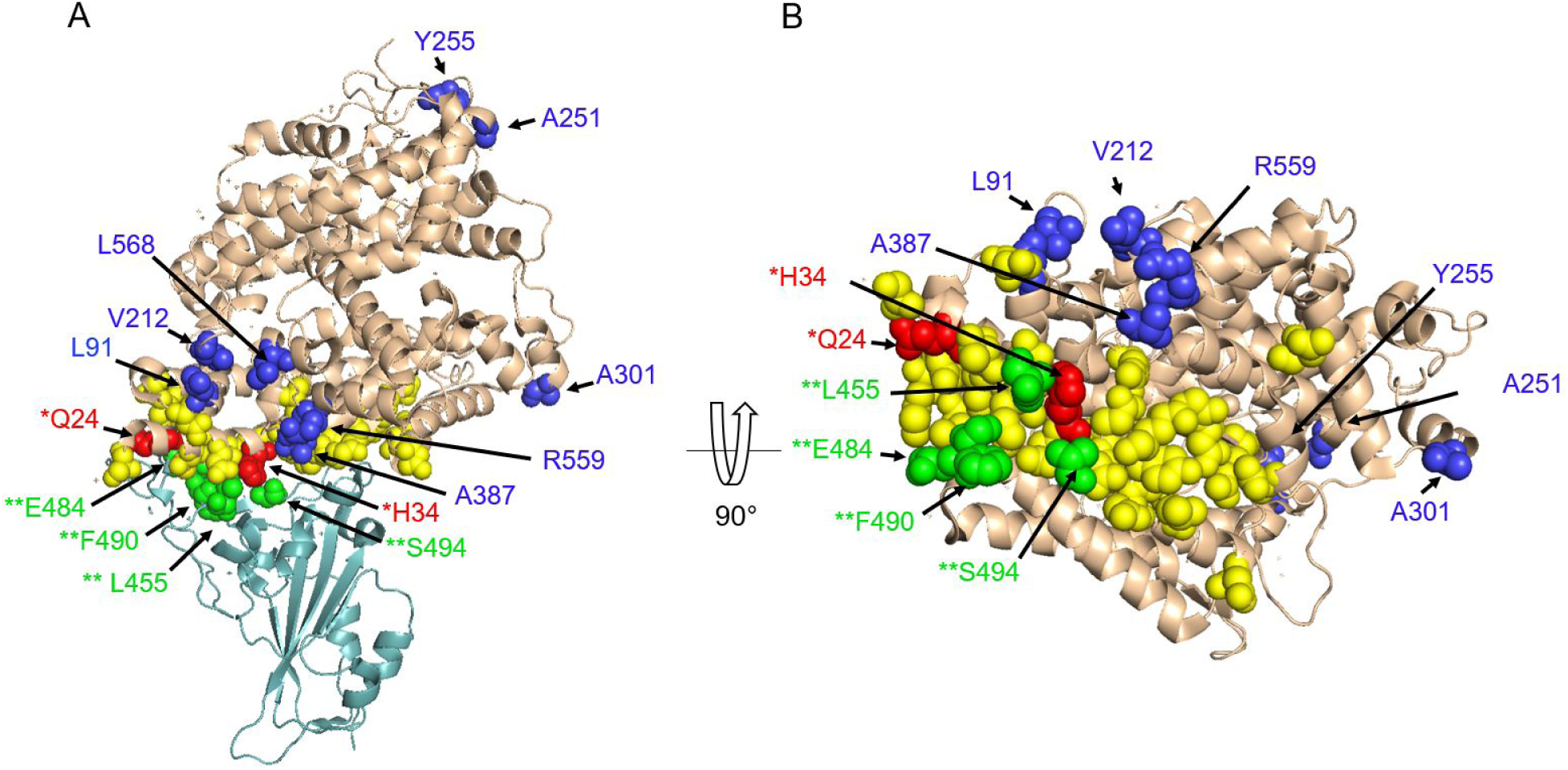
Residues under positive selection detected with CODEML and acceleration with phyloP in mammals. (**A**) ACE2 is represented in wheat cartoon with residues involved in the binding interface shown in yellow spheres. Dark blue and red spheres indicate residues in ACE2 that are accelerated and under positive selection. Red spheres represent residues that overlap with positions in the binding interface and are labeled with (*). The spike RBD is shown in light teal cartoon. Green spheres indicate residues on the SARS-CoV-2 spike protein under positive selection and are labeled with (**). (**B**) 90 degree rotation of the ACE2 protein.

This pattern of acceleration and positive selection in *ACE2* also holds for individual mammalian lineages. Using CODEML, positive selection was detected within the orders Chiroptera (LRT=346.40, ⍵=3.44 p<0.001), Cetartiodactyla (LRT=92.86, ⍵=3.83, p<0.001), Carnivora (LRT=65.66, ⍵=2.27. p<0.001), Primates (LRT=72.33, ⍵=3.16, p<0.001) and Rodentia (LRT=91.26, ⍵=1.77, p<0.001). Overall, bats had more positively selected sites with significant BEB scores (29 sites in Chiroptera compared to 10, 8, 7 and 15 sites in Cetartiodactyla, Carnivora, Primates and Rodentia, respectively). Positive selection at key sites for the binding of ACE2 and SARS-CoV-2 was only found in the bat-specific alignment. PhyloP was used to assess shifts in evolutionary rate within mammalian lineages, for each assessing signal relative to a neutral model trained on species from the specified lineage (Dataset S4.6-11, Fig. S6). We discovered six important binding residues, five of which showed evidence for positive selection, that are accelerated in one or more of Chiroptera, Rodentia, or Carnivora, with G354 accelerated in all of these lineages (Dataset S4.12).

Given pervasive signatures of adaptive evolution in *ACE2* across mammals, we next sought to test if any mammalian lineages are evolving particularly rapidly compared to the others. CODEML branch-site tests identified positive selection in both the ancestral Chiroptera branch (1 amino acid, ⍵=26.7, LRT= 4.22, p=0.039) and ancestral Cetartiodactyla branch (2 amino acids, ⍵=10.38, LRT= 7.89, p=0.004, Dataset S4.3) using 64 mammals. These residues did not correspond to known viral binding sites. We found no evidence for lineage-specific positive selection in the ancestral primate, rodent or carnivore lineages. PhyloP identified lineage-specific acceleration in Chiroptera, Carnivora, Rodentia, Artiodactyla and Cetaceans relative to mammals (Dataset S4.13-17, Fig. S7). Bats have a particularly high level of accelerated evolution (18 codons; p<0.05). Of these accelerated residues, T27 and M82 are known to be important for binding SARS-CoV-2, with some bat subgroups having amino acids predicted to lead to less favorable binding of SARS-CoV-2 (Fig. S1, Fig. S8). Surprisingly, a residue that is conserved overall in our 410 species alignment and in the mammalian subset, Q728, is perfectly conserved in all 37 species of bats except for fruit bats (Pteropodidae), which have a substitution from Q to E. These results support the theory that *ACE2* is under lineage-specific selective pressures in bats relative to other mammals.

### Positive selection in SARS-CoV-2 S protein

Positive selection was found using CODEML at sites L455, E484, F490 and S494 in the SARS-CoV-2 S sequence (⍵=1.15, LRT=116.7, p<0.001); however this signal was not particularly high, possibly due to the small sample size (N=8). All of these sites lie within or near the ACE2 SARS-CoV-2 S RBD binding sites (Fig. 3) (38).

## Discussion

Phylogenetic analysis of coronaviruses has demonstrated that SARS-CoV-2 most likely originated in a bat species (1). However, whether SARS-CoV-2 or the progenitor of this virus was transmitted directly to humans or through an intermediary host is not yet resolved. To determine if amino acid substitution analysis and structural information could be used to identify candidate intermediate host species, we undertook a deep comparative genomic, evolutionary and structural analysis of ACE2, the SARS-CoV-2 receptor in humans. To accomplish this we drew on the rapidly growing dataset of annotated vertebrate genomes as well as predicted protein sequences from recently acquired whole genome sequences produced by the Genomes 10K-affiliated Bat1K Consortium, Zoonomia, and Vertebrate Genomes Project, and other sources (39, 40). We conducted a phylogenetic analysis of ACE2 orthologs from 410 vertebrate species and made predictions of their likelihood to bind the SARS-CoV-2 S using a score that was based on amino acid substitutions at 25 consensus human ACE2 binding residues (13, 21). We supported these predictions with comprehensive homology modeling of the ACE2 binding site. We also tested the hypothesis that the ACE2 receptor is under selective constraints in different mammalian lineages, and correlated these results with data on the known species distribution of coronaviruses.

Several recent studies examined the role of ACE2 in SARS-CoV-2 binding and cellular infection, and its relationship to experimental and natural infections in different species (30, 41–46). Our study design differs substantially from those studies in several aspects: 1) we analyzed a larger number of primates, carnivores, rodents, cetartiodactyls and other mammalian orders, and an extensive phylogenetic sampling of fishes, birds, amphibians and reptiles; 2) we analyzed the full complement of S-binding residues across the ACE2 binding site, which was based on a consensus set from two independent studies (13, 21); 3) we used different methodologies to assess ACE2 binding capacity for SARS-CoV-2 S; and, 4) our study tested for selection and accelerated evolution across the entire ACE2 protein. While our results are strongly consistent with the results and conclusions of Melin and colleagues (44) on the predicted susceptibility of primates to SARS-CoV-2, particularly Old World primates, our work made predictions for a larger number of primates (N=39 vs N=27), bats (N=37 vs N=7), other mammals (N=176 vs N=5) and other vertebrates (N=158 vs N=0). When ACE2 from species in our study were compared with results of other studies there were many consistencies, such as for rodents, but some predictions that differ, such as the relatively high risk described for SARS-CoV-2 binding in pangolin and horse (45), civet (46), *Rhinolophus sinicus* bats (46) and turtles (45). In one recent study, binding affinity of soluble ACE2 for the SARS-CoV-2 S RBD was analyzed by saturation mutagenesis (26). Results obtained at each ACE2 binding residue were generally consistent with ours, particularly in the binding hotspot region of ACE2 residues 353-357. Importantly, as compared with other studies, our results greatly expanded the potential number of intermediate hosts and identified many more threatened species that could be infected by SARS-CoV-2 via their ACE2 receptors.

### Evolution of *ACE2*

Variation of *ACE2* in the human population is rare (17). We examined a large set of *ACE2* variants for their potential differences in binding to SARS-CoV-2 S and their relationship to selected and accelerated sites. We found rare variants that would result in missense mutations in 7 out of the 25 binding residues (Dataset S2). Some of those (e.g. E35K with an AF of 0.00001636) could reduce the virus binding affinity, thus potentially lowering the susceptibility to the virus in a very small fraction of the population. The analysis suggests that some variants (e.g. D38E) might not affect the binding while others (e.g. S19P) have uncertain effects. Further studies are needed to confirm and correctly address recent discoveries (18, 27, 28) and the data presented here, investigating the possible effect of these rare variants in specific populations.

When exploring patterns of codon evolution in ACE2, we found that a number of sites are evolving at different rates in the different lineages represented in our 410-species vertebrate alignment. Multiple ACE2 RBD residues important for the binding of SARS-CoV-2 are evolving rapidly across mammals, with two (Q24 and H34) under positive selection (Fig. 3, Fig. S5). Relative to other lineages analyzed, Chiroptera has a greater proportion of accelerated versus conserved residues, particularly at the SARS-CoV-2 S RBD, suggesting the possibility of selective forces on these codons in Chiroptera driven by their interactions with SARS-CoV-2-like viruses (Dataset S4.12, Fig. S8). Indeed, distinct signatures of positive selection found in bats and in the SARS-CoV S protein support this hypothesis that bats are evolving to tolerate SARS-CoV-2-like viruses.

### Relationship of the ACE2 binding score to known infectivity of SARS-CoV-2

Data on susceptibility of wild animals to SARS-CoV-2 is still very limited. It has been reported that a captive Malayan tiger was infected by SARS-CoV-2 (9) and that domestic cats, ferrets (47), rhesus macaques (48) and Syrian golden hamsters (49) are susceptible to experimental infection by SARS-CoV-2. These results agree with our predictions of ACE2 binding ability to SARS-CoV-2 S (Fig. 1, Dataset S1); 4/5 five species with demonstrated susceptibility to SARS-CoV-2 score *very high* (Rhesus macaque) or *medium* (domestic cat, tiger and Golden hamster). The only inconsistency was observed for ferrets, which had a *low* ACE2 binding score. This inconsistency could be related to the high infectivity dose used for experimental infection that likely does not correspond to virus exposure in nature. Dogs have low susceptibility to SARS-CoV-2 under experimental conditions (47), and score *low* for binding of their ACE2 to SARS-CoV-2 S. However, kidney cell lines derived from dog showed ACE2-dependent SARS-CoV-2 S entry, suggesting that *in vitro* experiments may be overestimating true infectivity potential (39, 50). Pigs (*low*), ducks (*very low*) and chickens (*very low*) were similarly exposed to SARS-CoV-2 and showed no susceptibility (47), providing further support of our methodology. A recent publication reporting that SARS-CoV-2 could use pig, masked palm civet and Chinese rufous horseshoe bat ACE2 expressed in HeLa cells were inconsistent with our predictions, while data for mouse was in agreement (1). Indeed, while mouse ACE2 scored *very low* in our analysis, pig and Chinese rufous horseshoe bat score *low*, while the masked palm civet scored *very low*. As for the ferret, high-level exposure to the virus *in vitro* could potentially result in infection via low affinity interactions with ACE2. Another possibility is that other cellular machinery present in the human HeLa cells is facilitating the infection, and that infectivity does not relate directly to ACE2 differences in these species. Confirmation of *in vitro* and *in vivo* susceptibility of these species under physiological conditions and with proper controls is clearly necessary. In addition, the expression of ACE2 varies across animal age, cell types, tissues and species (51, 52), which may lead to discrepancies between SARS-CoV-2 susceptibility gleaned from experimental infections or laboratory experiments and predictions made from the ACE2-based binding score.

### Mammals with high predicted risk of SARS-CoV-2 infection

Of the 19 catarrhine primates analyzed, 18/19 scored *very high* for binding of their ACE2 to SARS-CoV-2 S and one scored *high* (the Angola colobus); the 18 species scoring *very high* had 25/25 identical binding residues to human ACE2, including rhesus macaques (*Macaca mulatta*), which are known to be infected by SARS-CoV-2 and develop COVID-19-like clinical symptoms (3, 48). Our analysis predicts that all Old World primates are susceptible to infection by SARS-CoV-2 via their ACE2 receptors. Thus, many of the 21 primate species native to China could be a potential reservoir for SARS-CoV-2. The remaining primate species were scored as *high* or *medium*, with only the Gray mouse lemur and the Philippine tarsier scoring as *low*.

We were surprised to find that all three species of Cervid deer and 12/14 cetacean species have *high* scores for binding of their ACE2s to SARS-CoV-2 S. There are 18 species of Cervid deer found in China. Therefore, Cervid deer cannot be ruled out as an intermediate host for SARS-CoV-2. While coronavirus sequences have been found in white tailed deer (53) and gammacoronaviruses have been found in beluga whales (54, 55) and bottlenose dolphins (56) and are associated with respiratory diseases, the cellular receptor used by these viruses is not known.

### Other artiodactyls

A relatively large fraction (21/30) of artiodactyl mammals were classified with *medium* score for ACE2 binding to SARS-CoV-2 S. These include many species that are commonly found in Hubei Province and around the world, such as domesticated cattle, sheep and goats, as well as many species commonly found in zoos and wildlife parks (e.g., Masai giraffe, okapi, hippopotamus, water buffalo, scimitar horned oryx, and Dama gazelle). Although cattle MDBK cells were shown in one study to be resistant to SARS-CoV-2 *in vitro* (50), we propose immediate surveillance of common artiodactyl species for SARS-CoV-2 and studies of cellular infectivity, given our predictions. If ruminant artiodactyls can serve as a reservoir for SARS-CoV-2, it would have significant epidemiological implications as well as implications for food production and wildlife management (see below). It is noteworthy that camels and pigs, known for their ability to be infected by coronaviruses (35), both score *low* in our analysis. These data are consistent with results (discussed above) indicating that pigs cannot be infected with SARS-CoV-2 both *in vivo* (47) and *in vitro* (50).

### Rodents

Among the rodents, 7/46 species score *high* for ACE2 binding to SARS-CoV-2 S, with the remaining 11, 10 and 18 scoring *medium*, *low* or *very low*, respectively. Brown rats *(Rattus norvegicus*) and the house mouse (*Mus musculus*), scored *very low*, consistent with infectivity studies (1, 50). Given that wild rodent species likely come in contact with bats as well as with other predicted high risk species, we urge surveillance of *high* and *medium* binding likelihood rodents for the presence of SARS-CoV-2.

### Bats and other species of interest

Chiroptera (bats) represent a clade of mammals that are of high interest in COVID-19 research because several bat species are known to harbor coronaviruses, including those most closely related to the betacoronavirus SARS-CoV-2 (1). We analyzed ACE2 from 37 bat species of which 8 and 29 scored *low* and *very low*, respectively. These results were unexpected because the three *Rhinolophus* spp. including the Chinese rufous horseshoe bat are major suspects in the transmission of SARS-CoV-2, or a closely related virus, to humans (1). Globally, bats have been shown to harbour the highest diversity of betacoronaviruses in mammals tested (35) and show little pathology carrying these viruses (57). We found evidence for accelerated evolution at six RBD binding domain residues within the bat lineage, which is more than in any other lineage tested. Bats also had far more sites showing evidence of positive selection, including four binding domain residues, compared to other mammalian orders. This suggests that the diversity observed in bat ACE2 sequences may be driven by selective pressure from coronaviruses. Our results suggest that SARS-CoV-2 is not likely to use the ACE2 receptor in bats, which challenges a recent study showing that SARS-CoV-2 can infect HeLa cells expressing *Rhinolophus sinicus* ACE2 (1). If bats can be infected with SARS-CoV-2, the virus likely uses a different receptor. For example, the MERS-CoV, a betacoronavirus, uses CD26/DPP4 (16) while the porcine transmissible enteritis virus, an alphacoronavirus uses aminopeptidase N (ANPEP) (58). As detailed above, further *in vitro* and *in vivo* infectivity studies are required to fully understand the mode of transmission of susceptibility of bats to SARS-CoV-2.

### Carnivores

Recent reports of a Malayan tiger and a domestic cat infected by SARS-CoV-2 suggest that the virus can be transmitted to other felids (9, 47). Our results are consistent with these studies; 9/9 felids we analyzed scored *medium* for ACE2 binding of SARS-CoV-2 S. However, the masked palm civet (*Paguma larvata*), a member of the Viverridae family that is related to but distinct from Felidae, scored as *very low*. These results are inconsistent with transfection studies using civet ACE2 receptors expressed in HeLa cells (1), although these experiments have limitations as discussed above. While carnivores closely related to dogs (dingos, wolves and foxes) all scored *low*, experimental data supporting infection in dogs were inconsistent (47, 50, 59) so no conclusions can be drawn.

### Pangolins

Considerable controversy surrounds reports that pangolins can serve as an intermediate host for SARS-CoV-2. Pangolins were proposed as a possible intermediate host (22) and have been shown to harbor related coronaviruses. In our study, ACE2 of Chinese pangolin (*Manis pentadactyla*), Sunda pangolin (*Manis javanica*), and white bellied pangolin (*Phataginus tricuspis*) had *low* or *very low* binding score for SARS-CoV-2 S. Neither experimental infection nor *in vitro* infection with SARS-CoV-2 has been reported for pangolins. As for ferrets and bats, if SARS-CoV-2 infects pangolins it may be using a receptor other than ACE2, based on our analysis.

### Other vertebrates

Our analysis of 29 orders of fishes, 29 orders of birds, 3 orders of reptiles and 2 orders of amphibians predicts that the ACE2 proteins of species within these vertebrate classes are not likely to bind SARS-CoV-2 S. Thus, vertebrate classes other than mammals are not likely to be an intermediate host or reservoir for the virus, despite predictions reported in a recent study (45), unless SARS-CoV-2 can use another receptor for infection. With many different non-mammal vertebrates sold in the seafood and wildlife markets of Asia and elsewhere, it is still important to determine if SARS-CoV-2 can be found in non-mammalian vertebrates.

### Relevance to Threatened Species

Among the 103 species that scored *very high*, *high* and *medium* for ACE2 SARS-CoV-2 S RBD binding, 41 (40%) are classified in one of three ‘Threatened’ categories (*Vulnerable*, *Endangered*, and *Critically Endangered*) on the IUCN Red List of Threatened Species, five are classified as *Near Threatened*, and two species are classified as *Extinct in the Wild* (Dataset S1)(60). This represents only a small fraction of the threatened species potentially susceptible to SARS-CoV-2. For example, all 20 catarrhine primate species in our analysis, representing three families (Cercopithecidae, Hylobatidae, and Hominidae) scored *very high*, suggesting that all 185 species of catarrhine primates, most of which are classified Threatened (62), are potentially susceptible to SARS-CoV-2. Similarly, all three species of deer, representatives of a family of ~92 species (Cervidae), scored as *high* risk, as did species representing Cetacea (baleen and toothed whales), and both groups contain a number of threatened species. Toothed whales have potential for viral outbreaks and have lost function of a gene key to the antiviral response in other mammalian lineages (61). If they are susceptible to SARS-CoV-2, human-to-animal transmission could pose a risk through sewage outfall (62) and contaminated refuse from cities, commercial vessels and cruise liners (63). In contrast, some threatened species scored *low* or *very low*, such as the giant panda (*low*), potentially positive news for these at risk populations.

Our results have practical implications for populations of threatened species in the wild and those under human care (including those in zoos). Established guidelines for minimizing potential human to animal transmission should be implemented and strictly followed. Guidelines for field researchers working on great apes established by the IUCN have been in place since 2015 in response to previous human disease outbreaks (64) and have received renewed attention because of SARS-CoV-2 (64–66). For zoos, guidelines in response to SARS-CoV-2 have been distributed by several Taxon Advisory Groups of the North American Association of Zoos and Aquariums (AZA), the American Association of Zoo Veterinarians (AAZV), and the European Association of Zoo and Wildlife Veterinarians (EAZWV), and these organizations are actively monitoring and updating knowledge of species in human care considered to be potentially sensitive to infection (67, 68). Although *in silico* studies suggest potential susceptibility of diverse species, verification of infection potential is warranted, using cell cultures, stem cells, organoids, and other methods that do not require direct animal infection studies. Zoos and other facilities that maintain living animal collections are in a position to provide such samples for generating crucial research resources by banking tissues, and cryobanking viable cell cultures in support of these efforts.

### Animal models for COVID-19

A variety of animal models have been developed for studying SARS and MERS coronavirus infections (69). Presently, there is a tremendous need for animal models for studying SARS-CoV-2 infection and pathogenesis, as the only species currently known to be infected and show similar symptoms of COVID-19 is rhesus macaque. Non-human primate models have proven to be highly valuable for other infectious diseases, but are expensive to maintain and numbers of experimental animals are limited. Our results provide an extended list of potential species that might be useful as animal models for SARS-CoV-2 infection and pathogenesis, including Chinese hamster and Syrian/Golden hamster (49), and large animals maintained for biomedical and agricultural research (e.g., domesticated sheep and cattle).

### Conclusions

We predict that species scored as *very high* and *high* for SARS-CoV-2 S binding to ACE2 will have a high probability of becoming infected by the virus. We also predict that many species having a *medium* score have some risk of infection, and species scored as *very low* and *low* are unlikely to be infected by SARS-CoV-2 via the ACE2 receptor. Importantly, our predictions are based solely on *in silico* analyses and must be confirmed by direct experimental data. Until such time, other than for species in which SARS-CoV-2 infection has been demonstrated to occur using ACE2, we urge caution not to over-interpret the predictions made in the present study. This is especially important with regards to species, endangered or otherwise, in human care. While species ranked *high* or *medium* may be susceptible to infection based on the features of their ACE2 residues, pathological outcomes may be very different among species depending on other mechanisms that could affect virus replication and spread to target cells, tissues, and organs within the host. Furthermore, we cannot exclude the possibility that infection in any species occurs via another cellular receptor, as has been shown for other betacoronaviruses. Nonetheless, our predictions provide a useful starting point for selection of appropriate animal models for COVID-19 research and for identification of species that may be at risk for human-to-animal or animal-to-animal transmissions by SARS-CoV-2. The approach we used for ACE2 can be extended to other cellular proteins known to be involved in coronavirus infection and immunity to better understand infection, transmission, inflammatory responses and disease progression.

## Materials and Methods

### Angiotensin I converting enzyme 2 (ACE2) coding and protein sequences

All human ACE2 orthologs for vertebrate species, and their respective coding sequences, were retrieved from NCBI Protein (March 20, 2020) (70). ACE2 coding DNA sequences were extracted from available or recently sequenced unpublished genome assemblies for 123 other mammalian species, with the help of genome alignments and the human or within-family ACE2 orthologs. The protein sequences were predicted using AUGUSTUS v3.3.2 (71) or CESAR v2.0 (72) and the translated protein sequences were checked against the human ACE2 orthologue. ACE2 gene predictions were inspected and manually curated if necessary. For four bat species (*Micronycteris hirsuta*, *Mormoops blainvillei*, *Tadarida brasiliensis* and *Pteronotus parnellii*) the ACE2 coding region was split into two scaffolds which were merged, and for *Eonycteris spelaea* a putative 1bp frameshift base error was corrected. Eighty ACE2 predictions were obtained from the Zoonomia project, 19 from the Hiller Lab, 12 from the Koepfli lab, 8 from the Lewin lab and 4 from the Zhou lab. The source, and accession numbers for the genomes or proteins retrieved from NCBI are listed in Dataset S1. The final set of ACE2 sequences comprises 410 vertebrate species. To assure alignment robustness, the full set of coding and protein sequences were aligned independently using Clustal Omega (73), MUSCLE (74) and COBALT (75) all with default parameters. All resulting protein alignments were identical. Clustal Omega alignments were used in the subsequent analysis. Each amino acid replacement present in our dataset was classified as neutral, semi-conservative and non-conservative as in Clustal Omega.

### Identification of ACE2 residues involved in binding to SARS-CoV-2 S protein

We identified 22 ACE2 protein residues that were previously reported to be critical for the effective binding of ACE2 RBD and SARS-CoV-2 S (13, 21). These residues include S19, Q24, T27, F28, D30, K31, H34, E35, E37, D38, Y41, Q42, L45, L79, M82, Y83, N330, K353, G354, D355, R357, and R393. All these residues were identified from the co-crystallization and structural determination of SARS-CoV-2 S and ACE2 RBD (13, 21). The known human ACE2 RBD glycosylation sites N53, N90 and N322 were also included in the analyzed residue set (11).

### ACE2 and SARS-CoV-2 binding ability prediction

Based on the known interactions of ACE2 and SARS-CoV-2 residues, we developed a set of rules for predicting the likelihood of the SARS-CoV-2 S binding to ACE2. Each species was classified in one of five categories: *very high*, *high*, *medium*, *low* or *very low* likelihood of binding SARS-CoV-2 S. Species in the *very high* category have at least 23/25 critical residues identical to the human; have K353, K31, E35, M82, N53, N90 and N322; do not have N79; and have only conservative substitutions among the non-identical 2/25 residues. Species in the *high* group have at least 20/25 residues identical to the human; have K353; have only conservative substitutions at K31 and E35; do not have N79; and can only have one non-conservative substitution among the 5/25 non-identical residues. Species scoring *medium* have at least 20/25 residues identical to the human; can only have conservative substitutions at K353, K31, and E35; and can have up to two non-conservative substitutions in the 5/25 non-identical residues. Species in the *low* category have at least 18/25 residues identical to the human; can only have conservative substitutions at K353; can have up to three non-conservative substitutions on the remaining 7/25 non-identical residues. Lastly, species in the *very low* group have less than 18/25 residues identical to the human or have at least four non-conservative substitutions in the non-identical residues.

### Protein structure analysis

We applied an orthogonal approach to assess the likelihood of binding of a sampling of species that were predicted to bind SARS-CoV-2 across the categories of *high*, *medium*, *low* or *very low* likelihood of binding. ACE2 amino acid sequences from 28 species were extracted from the multiway alignment and loaded into SWISS-MODEL (25) in order to generate homology derived models. The output files were aligned to the crystal structure 6MOJ (13) in order to assess the overall similarities to human ACE2. We used two recently solved crystal structures of the complex for ACE2 and SARS-CoV-2 S RBD, 6MOJ (13) and 6VW1 (21) as ground truth for the human ACE2/SARS-CoV-2 S interaction. In the program CHIMERA (76), we utilized the rotamer function to model each individual variant that species exhibit separately, and chose the rotamer with the least number of clashes, retaining the most initial hydrogen bonds and containing the highest probability of formation as calculated by CHIMERA from the Dunbrack 2010 backbone-dependent rotamer library (77). The rotamer was then evaluated in the context of its structural environment and assigned a score based on likelihood of interface disruption. Neutral (N) was assigned if the residue maintained a similar environment as the original residue, and was predicted to maintain or in some cases increase affinity. Weakened (W) was assigned if hydrophobic contacts were lost and contacts that appear disruptive are introduced that are not technically clashes. Unfavorable (U) was assigned if clashes are introduced and/or a hydrogen bond is broken. Additional structural visualizations were generated in Pymol (78).

### Human variants analysis

All variants at the 25 residues critical for effective SARS-CoV-2-ACE2 binding (11, 21, 79) were compiled from from dbSNP (31), 1KGP (32), Topmed (33), UK10K (34) and CHINAMAP (28). Specific population frequencies were obtained from gnomAD v.2.1.1 (17).

### Phylogenetic reconstruction of the vertebrate *ACE2* species tree

The multiple sequence alignment of 410 ACE2 orthologous protein sequences from mammals, birds, fishes, reptiles and amphibians was used to generate a gene tree using the maximum likelihood method of reconstruction, as implemented in IQTREE (80). The best fit model of sequence evolution was determined using ModelFinder (81) and used to generate the species phylogeny. A total of 1000 bootstrap replicates were used to determine node support using UFBoot (82).

### Identifying sites undergoing positive selection

Signatures of site-specific positive selection in the *ACE2* receptor were explored using CODEML, part of the Phylogenetic Analysis using Maximum Likelihood (PAML, (83)) suite of software. Given CODEML’s computational complexity, a smaller subset of mammalian taxa (N=64, Dataset S1), which included species from all prediction categories mentioned above, was used for selection analyses. To calculate likelihood-derived dN/dS rates (⍵), CODEML utilises both a species tree and a codon alignment. The species tree for all 64 taxa was calculated using IQTREE (80) and the inferred best-fit model of sequence evolution (JTT+F+R4). This gene topology was generally in agreement with the 410 taxa tree, however bats were now sister taxa to Perissodactyla. Therefore all selection analyses were run using both the inferred gene tree, and a modified tree with the position of bats manually modified to reflect the 410 taxa topology. All species trees used were unrooted. A codon alignment of the 64 mammals was generated using pal2nal (84) with protein alignments generated with Clustal Omega (73) and their respective CDS sequences.

Site-models M7 (null model) and M8 (alternative model) were used to identify *ACE2* sites undergoing positive selection in mammals. Both M7 and M8 estimate ⍵ using a beta distribution and 10 rate categories per site with ⍵<=1 (neutral or purifying selection), but with an additional 11^th^ category allowing ⍵ >1 (positive selection) in M8. A likelihood ratio test (LRT) calculated as 2*(lnL_alt_ – lnL_null_), comparing the fit of both null and alternative model likelihoods was carried out, with a p-value calculated assuming a chi-squared distribution. Sites showing evidence of positive selection were identified by a significant (>0.95) Bayes Empirical Bayes (BEB) score, and validated by visual inspection of the protein alignment. To explore order-specific instances of positive selection, separate multiple sequence alignments and gene trees for Chiroptera (N=37), Cetartiodactyla (N=45), Carnivora (N=44), Rodentia (N=46) and Primates (N=39) were also generated and explored using M7 vs. M8 in CODEML.

In addition to site-models, branch-site model A1 (null model) and model A (alternative model) were also implemented targeting various mammalian orders, specifically Chiroptera, Cetartiodactyla, Rodentia and Primates, to identify lineage-specific positive selection in the *ACE2* receptor sequence. Branch-site Model A1 constrains both the target foreground branch (Carnivora, Chiroptera, Cetartiodactyla, Rodentia and Primates) and background branches to ⍵<=1, while the alternative Model A allows positive selection to occur in the foreground branch. Null and alternative models were compared using LRTs as above, with significant BEB sites identified.

We also looked for positively selected sites in the viral spike protein, using SARS-CoV-2 (MN908947.3), Bat coronavirus RaTg13 (MN996532.1), Bat SARS-like coronavirus isolate Rs4231 (KY417146.1), SARS-related coronavirus strain BtKY72 (KY352407.1), SARS coronavirus Urbani (AY278741.1), SARS coronavirus PC4-227 (AY613950.1), Coronavirus BtRs-BetaCoV/YN2018B (MK211376.1) and the more divergent Bat Hp-betacoronavirus/Zhejiang2013 (NC_025217.1) viral strains. Protein and codon alignments were generated as above, with the viral species tree inferred using full genome alignments of all strains generated with Clustal Omega (73). Site-test models were applied using CODEML, and significant BEB sites identified.

### Analysis for departure from neutral evolutionary rate in ACE2 with PHAST

Neutral models were trained on the specified species sets (Dataset S4) using the REV nucleotide substitution model implemented in phyloFit using an expectation maximization algorithm for parameter optimization. The neutral model fit was based on third codon positions to approximate the neutral evolution rate specific to the *ACE2* gene, using a 410-species phylogenetic tree generated by IQTREE as described above and rooted on fishes. The program phyloP was then used to identify codons undergoing accelerated or conserved evolution relative to the neutral model using --features to specify codons, --method LRT --mode CONACC, and --subtree for lineage-specific tests, with p-values thus assigned per codon based on a likelihood ratio test. P-values were corrected for multiple testing using the Benjamini-Hochberg method (36) and sites with a corrected p-value less than 0.05 were considered significant. PhyloFit and phyloP are both part of the PHAST package v1.4 (85, 86).

## Acknowledgements

We thank Lawrence Stern for helpful discussions on homology modeling. We thank Pavel Dobrynin, Paul Frandsen, Taylor Hains, Sergei Kliver, and Alice Mouton for extracting and contributing ACE2 sequences from unpublished genomes. We thank Shirley Xue Li and Kate Megquier for help in data compilation. We thank Pierre Comizzoli, Budhan Pukazhenthi, and Nucharin Songasasen for valuable comments that improved the manuscript. This work was supported by the Robert and Rosabel Osborne Endowment (HAL). KLT is the recipient of a Distinguished Professor award from the Swedish Research Council. ECT is funded by an Irish Research Council Laureate Award. KCK is supported by a UCSF Discovery Fellowship and the Gladstone Institutes. GMH is funded by an Ad Astra Fellowship at University College Dublin. EKK, DPG and RS were supported by NIH R01HG008742. The research conducted in this study was coordinated as part of the Genome 10K Consortium, which includes the Bat1K, Zoonomia, the Vertebrate Genomes Project, and the Earth BioGenome Project.

