## Supplementary Information for "Broad Host Range of SARS-CoV-2 Predicted by Comparative and Structural Analysis of ACE2 in Vertebrates"

**This PDF file includes:**

Supplementary text
Figures S1 to S8
Legends for Datasets S1 to S4
SI References

**Other supplementary materials for this manuscript include the following:**

Datasets S1 to S4

**Supplementary Information Text**

Comparison of protein structure analysis methodologies. Of the 16 residues in the structural homology assessment performed in this study (Fig. S1), just one *unfavorable* substitution, K353H, differentiated mouse from the four other species tested in the Zhou and collaborators study (human, pig, Masked palm civet and Chinese rufous horseshoe bat)(1). Mouse was the single species whose ACE2 did not bind SARS-CoV-2 in that study (1). The K353H substitution has been shown experimentally to abrogate binding of ACE2 to SARS-CoV spike protein (2). We used this site to test another recently published approach that assessed binding affinity between ACE2 and SARS-CoV-2 using theoretical changes in Gibb's free energy (3). We estimated the change in Gibbs free energy for the K353H substitution for both SARS-CoV, using the template PDB 2AJF as well as the SARS-CoV-2 using the PDB 6M0J using the SSIPe program (4). The output suggested that the $\Delta\Delta G_{bind}$ in the SARS-CoV-2 would be 0.460 kcal/mol and in the SARs-CoV 0.594 kcal/mol, neither of which would be predicted to cause a disruption in affinity significant enough to abrogate binding (3, 4). Therefore, despite offering the potential to quantify predicted binding affinities, without information from experimentally derived binding assays, the Gibb's free energy method may not accurately represent the impact of different substitutions along the binding interface.

Another recent body of work emerged while this manuscript was under preparation, which provided a wealth of experimentally derived insights into the impact on binding to SARS-CoV-2 Spike protein when each residue of the ACE2 binding interface was substituted for each amino acid (5). We compared the predictions that we generated for residues that varied along the binding interface from 28 species with the results obtained by Procko, to determine the percent discordance and concordance between approaches (Table S1). Discordant values were calculated as 16.6% of the 55 substitutions analyzed using the following rules: discordant substitutions are designated as *neutral* (N) from this analysis and (--) from the Procko analysis; *unfavorable* (U) from this analysis and (0, +, ++) from the Procko analysis; *weaken* (W) from this analysis and (+, ++) from the Procko analysis according to Table S1. Residues that were calculated to be discordant between these two analyses were indicated by an (*) in Fig. S1.

| | Species | | Q24 | T27 | D30 | K31 | H34 | E35 | E37 | D38 | Y41 | Q42 | L45 | L79 | M82 | Y83 | K353 | G354 |
|---|---|---|---|---|---|---|---|---|---|---|---|---|---|---|---|---|---|---|
| High | Odocoileus virginianus texanus | *White-tailed deer* | . . | . . | E N | . . | . . | . . | . . | . . | . . | . . | . . | M N | T N | . . | . . | . . |
| | Rangifer tarandus | *Reindeer* | . . | . . | E N | . . | . . | . . | . . | . . | . . | . . | . . | M N | T N | . . | . . | . . |
| | Eulemur flavifrons | *Blue-eyed black lemur* | E N | A N | . . | . . | . . | . . | . . | . . | . . | . . | . . | . . | T N | . . | . . | . . |
| | Propithecus coquereli | *Coquerel's sifaka* | . . | . . | . . | . . | . . | . . | . . | . . | . . | . . | . . | . . | T N | . . | . . | . . |
| | Dipodomys stephensi | *Stephens's kangaroo rat* | L U | . . | . . | N W | Q N | . . | . . | . . | . . | . . | . . | . . | I N | . . | . . | . . |
| Medium | Bos taurus | *Cattle* | . . | . . | E N | . . | . . | . . | . . | . . | . . | . . | . . | M N | T N | . . | . . | . . |
| | Felis catus | *Cat* | L U | . . | E N | . . | . . | . . | . . | E N | . . | . . | . . | . . | T N | . . | . . | . . |
| | Panthera tigris altaica | *Siberian tiger* | L U | . . | E N | . . | . . | . . | . . | E N | . . | . . | . . | . . | T N | . . | . . | . . |
| | Mesocricetus auratus | *Golden hamster* | . . | . . | . . | . . | Q N | . . | . . | . . | . . | . . | . . | . . | N U* | . . | . . | . . |
| Low | Sus scrofa | *Pig* | L U | . . | E N | . . | L U | . . | . . | . . | . . | . . | . . | I N | T N | . . | . . | . . |
| | Ailuropoda melanoleuca | *Giant panda* | L U | . . | E N | . . | Y U | . . | . . | . . | . . | . . | . . | H W | T N | . . | . . | . . |
| | Canis lupus familiaris | *Dog* | L U | . . | E N | . . | Y U | . . | . . | E N | . . | . . | . . | . . | T N | . . | . . | . . |
| | Rhinolophus pearsonii | *Pearson's horseshoe bat* | . . | I N | . . | . . | R N | . . | . . | . . | H W | E W | . . | . . | D U | . . | . . | D U |
| | Dipodomys ordii | *Ord's kangaroo rat* | L U | . . | . . | N W | Q N | . . | . . | . . | . . | . . | . . | . . | I N | . . | . . | . . |
| Very Low | Catagonus wagneri | *Chacoan peccary* | L U | . . | E N | . . | L U | . . | . . | . . | . . | . . | . . | T W* | T N | . . | . . | . . |
| | Mustela putorius furo | *Ferret* | L U | . . | E N | . . | Y U | . . | . . | E N | . . | . . | . . | H W | T N | . . | . . | R U |
| | Paguma larvata | *Masked palm civet* | L U | . . | E N | T U | Y U | . . | Q N | E N | . . | . . | V N | . . | T N | . . | . . | . . |
| | Hipposideros armiger | *Great roundleaf bat* | L U | E U | . . | . . | T U* | . . | . . | . . | H W | L U* | . . | R W* | D U | . . | . . | . . |
| | Hipposideros galeritus | *Cantor's roundleaf bat* | S U | I N | . . | . . | T U* | D W* | . . | E N | H W | . . | . . | . . | D U | . . | . . | . . |
| | Hipposideros pratti | *Pratt's roundleaf bat* | L U | E U | . . | . . | T U* | . . | . . | . . | H W | L U* | . . | R W* | D U | . . | . . | . . |
| | Rhinolophus ferrumequinum | *Greater horseshoe bat* | L U | K U* | . . | D W | S W* | . . | . . | N N | H W | . . | . . | . . | N U* | F W | . . | . . |
| | Uropsilus gracilis | *Gracile shrew mole* | . . | E U | E N | N W | R N | N W | . . | N N | . . | K N/A | . . | I N | Q U | F W | M W | K U |
| | Manis javanica | *Sunda pangolin* | E N | . . | E N | . . | S W* | . . | . . | E N | . . | . . | . . | I N | N U* | . . | . . | H U |
| | Manis pentadactyla | *Chinese pangolin* | E N | . . | E N | . . | S W* | . . | . . | E N | . . | . . | . . | I N | N U* | . . | . . | H U |
| | Otolemur garnettii | *Northern greater galago* | . . | . . | . . | N W | R N | . . | . . | E N | H W | . . | . . | I N | T N | . . | . . | D U |
| | Mus musculus | *House mouse* | N U | . . | N N | . . | Q N | . . | . . | . . | . . | . . | . . | T W* | S U | F W | H U | . . |

* discordant with Procko et al

**Fig. S1.** Evaluation of binding contacts between host ACE2 and SARS-CoV-2 in 28 representative species selected from *very low*, *low*, *medium* and *high* binding score groups, and for each residue in the ACE2 binding interface that varied from human (55 substitutions in 16 residues). For each residue, amino acid substitutions are shown on the left as white boxes, with sites matching human ACE2 shown in gray. For each residue, the evaluation of the binding contact is shown on the right as *neutral* (N; blue box), w*eakening* (W; orange box); or *unfavorable* (U; red box), with sites matching human ACE2 in blue. Evaluations discordant with Procko (5) are marked with an asterix and lighter background color.

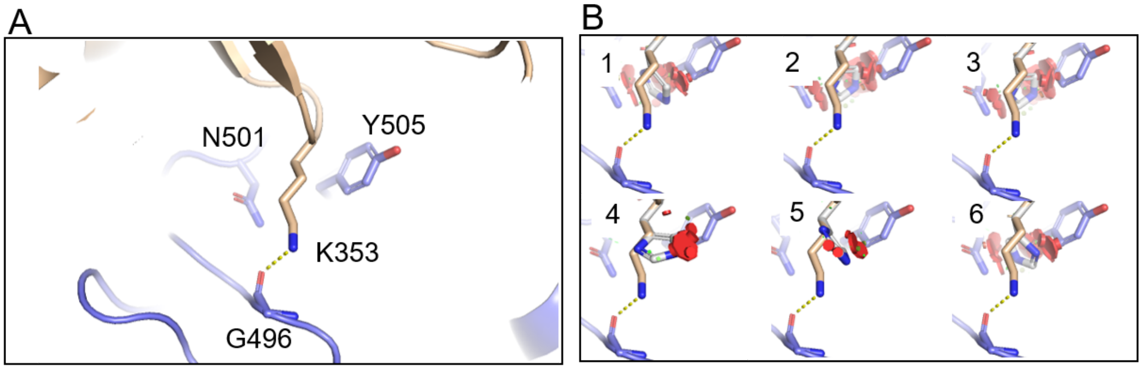


**Fig. S2.** Selecting the best-fit rotamer for K353H. (**A**) Cartoon representation of ACE2 K353 depicted in wheat stick with a dotted line representing the hydrogen bond to SARS-CoV-2 spike (S) G496. Y505 and N501 from the S RBD, which are contact residues for ACE2 K353 are shown. (**B**) All Dunbrack derived rotamers for histidine show clashes as red disks at this position, with rotamer 5 showing the least steric hindrance. In addition, substitution to histidine at this position would eliminate the hydrogen bond between the 353 position on ACE2 and the S RBD at G496. This substitution was designated as being potentially *unfavorable* due to the loss of the hydrogen bond in addition to the steric hindrance.

**Fig. S3 (separate file).** Phylogenetic tree of ACE2 proteins in mammals, rooted on fish.

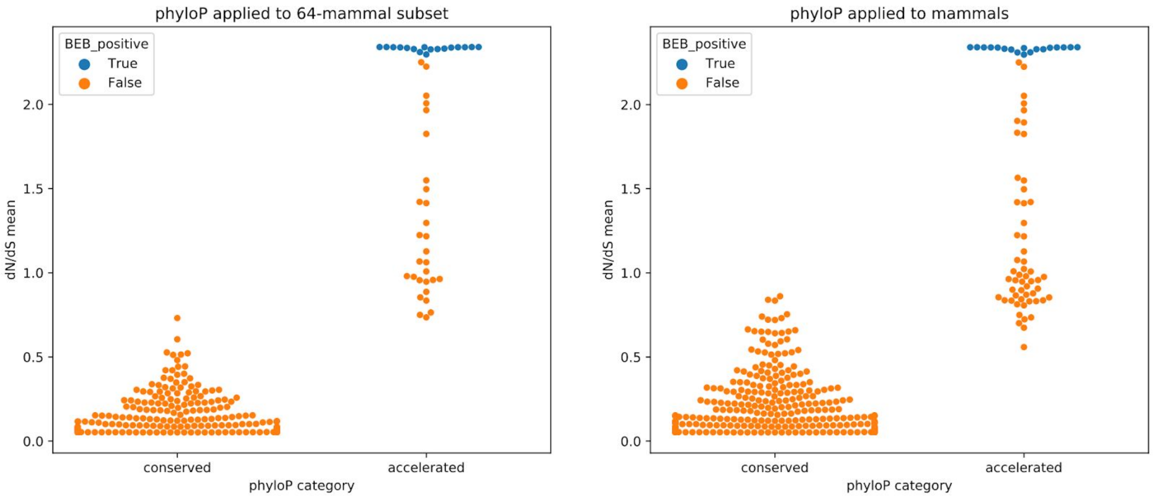


**Fig. S4.** Significant results from phyloP, both conserved and accelerated, for ACE2 codons compared with CODEML BEB scores. Left panel shows phyloP results for the 64-mammals subset used in the mammal CODEML analysis. Right panel shows phyloP results for all mammals in the alignment. The y-axis represents dN/dS values calculated by CODEML, x-axis indicates whether the codons were classified as conserved or accelerated by phyloP. All dots are significant results from phyloP, blue dots are also significantly positively selected from CODEML.

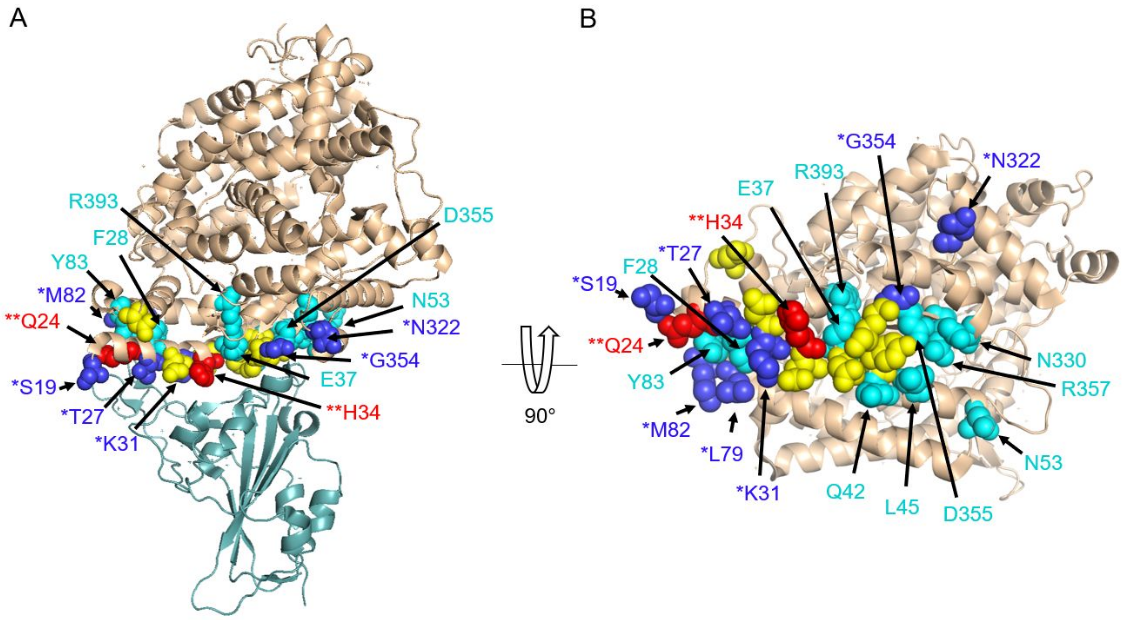


**Fig. S5.** Residues under accelerated evolution in mammals, overlapping the binding interface, as detected using phyloP. (**A**) The SARS-CoV-2 spike RBD is shown in light teal cartoon. ACE2 is shown in wheat cartoon with residues involved in the binding interface shown in yellow spheres. (*) Dark blue and red spheres indicate ACE2 residues that are accelerated, under positive selection and overlapping the binding interface. Cyan spheres indicate ACE2 residues that are conserved. (**) Red spheres also demonstrate positive selection with CODEML. (**B**) 90 degree rotation of the ACE2 protein.

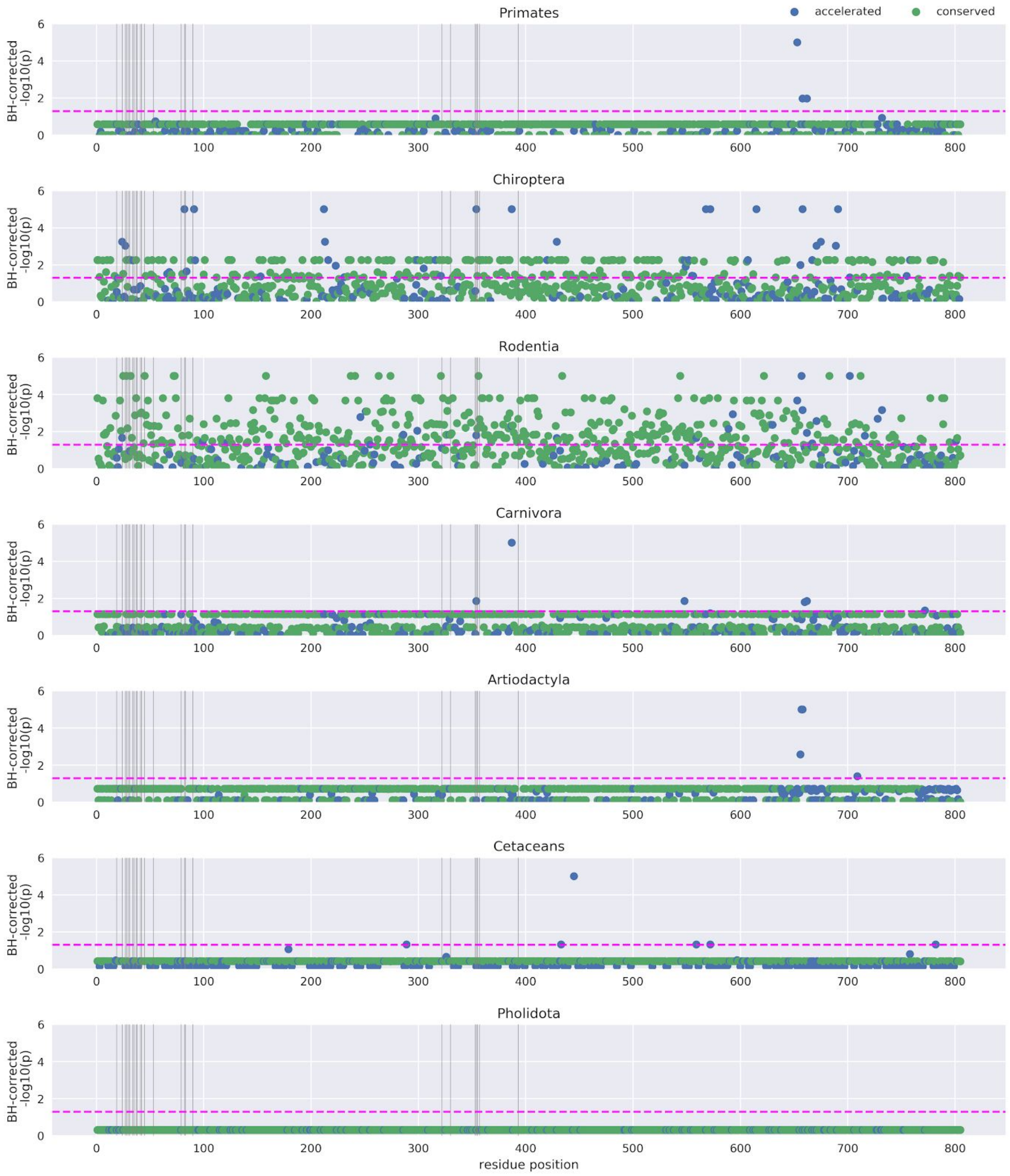


**Fig. S6.** Intralineage phyloP results for all ACE2 codons. PhyloP signal was assessed at all ACE2 codons for various mammalian lineages against neutral models trained on those lineages, thereby identifying intralineage signals of shifts in evolutionary rate. Green dots indicate codons classified as conserved and blue dots accelerated. Vertical grey lines indicate important binding residues in ACE2. The x-axis indicates the corresponding position in the ACE2 protein for each codon, and the y-axis indicates the phyloP p-value for each codon.

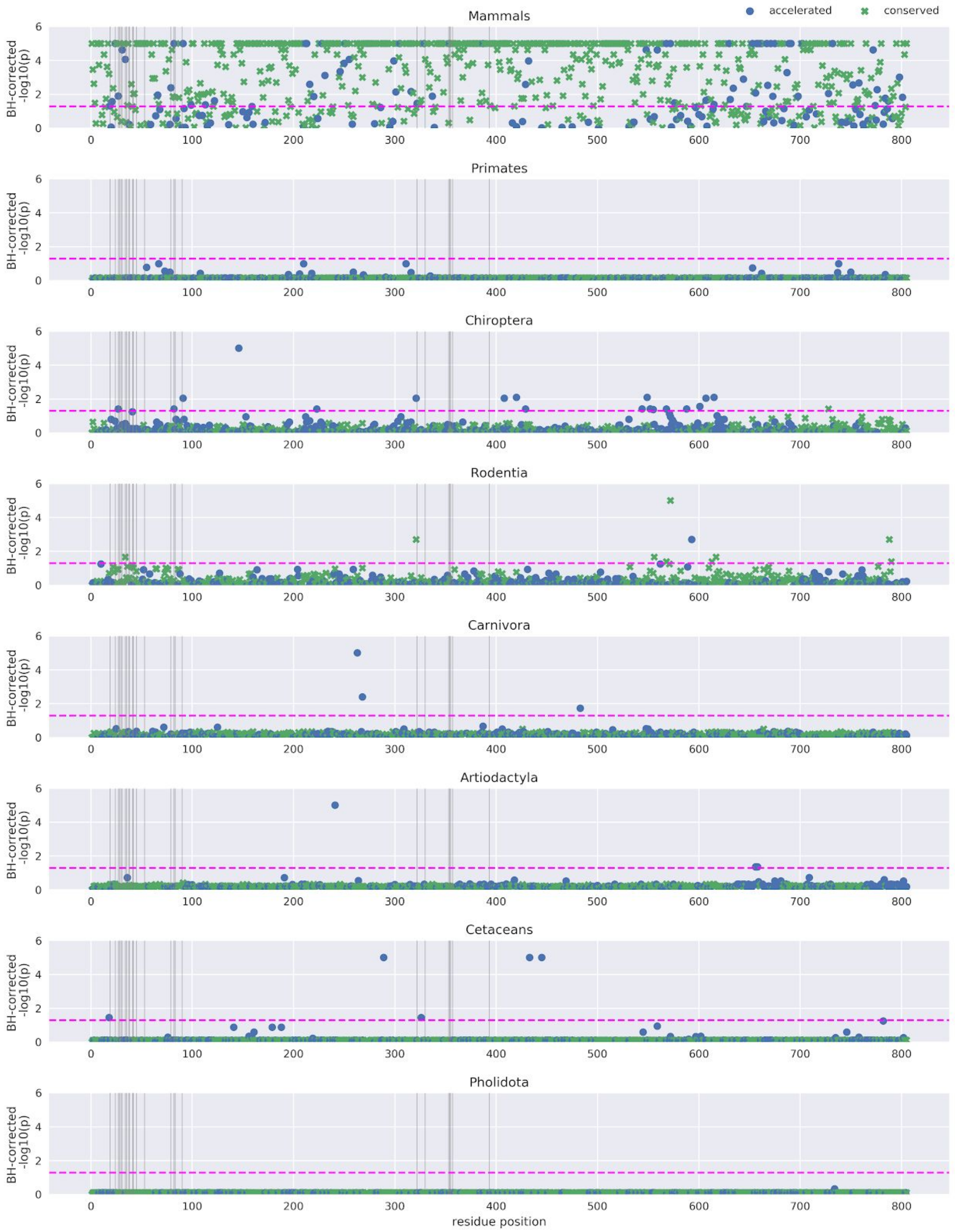


**Fig. S7.** PhyloP results for mammalian lineages against a mammal neutral model. PhyloP signal was assessed at all ACE2 codons for various mammalian lineages against a neutral model trained on all mammalian species in the alignment. Green dots indicate codons classified as conserved and blue dots accelerated. Vertical grey lines indicate important binding residues in ACE2. The x-axis indicates the corresponding position in the ACE2 protein for each codon, and the y-axis indicates the phyloP p-value for each codon.

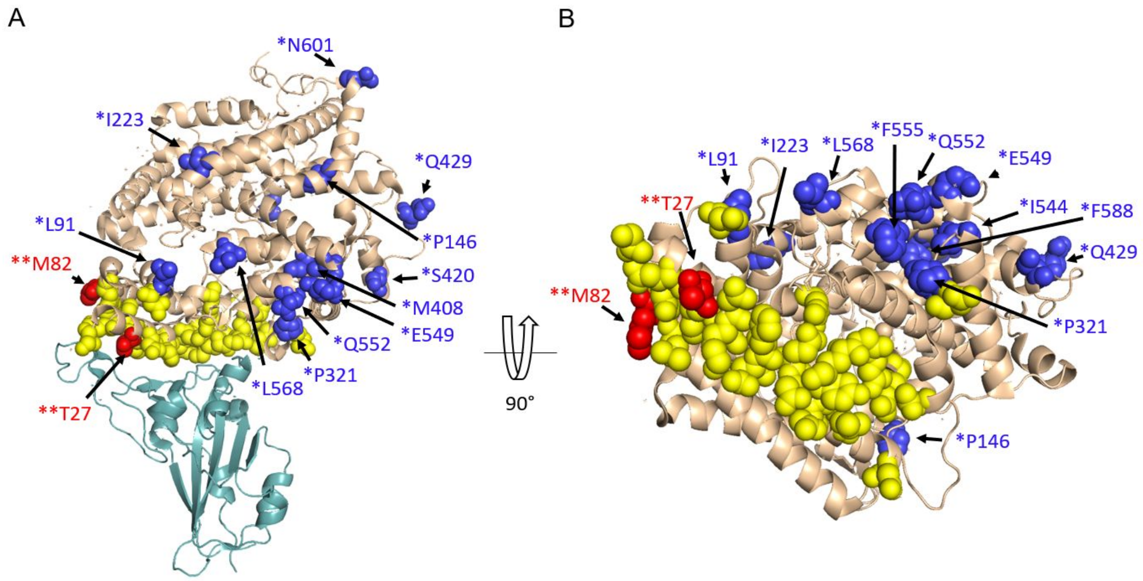


**Fig. S8.** Residues under acceleration with phyloP in chiroptera relative to mammals. (**A**) The SARS-CoV-2 spike RBD is shown in light teal cartoon. ACE2 is shown in wheat cartoon with residues involved in the binding interface shown in yellow spheres. Dark blue and red spheres indicate residues that are accelerated in bats relative to mammals. Red spheres also overlap the binding interface. (**B**) 90 degree rotation of the ACE2 protein.

**Table S1.** Comparing structure-based predictions and experimentally derived values for the impact of substitutions along the ACE2/SARS-CoV-2 spike RBD interface.

| Current work | Procko | Number matched |
|---|---|---|
| N | - | 13 |
| N | + | 1 |
| N | ++ | 3 |
| N | 0 | 1 |
| U | 1 | 12 |
| U | -- | 7 |
| U | ++ | 2 |
| U | 0 | 2 |
| W | - | 8 |
| W | + | 4 |
| W | ++ | 1 |
| W | 0 | 1 |

**Dataset S1 (separate file).** Variation at 25 residues critical for ACE2 and SARS-CoV-2 binding in 410 vertebrate species.

**Dataset S2 (separate file).** Table showing the human variant analysis on the 25 critical ACE2 binding residues.

**Dataset S3 (separate file).** Phylogenetic tree of ACE2 proteins in 410 vertebrate species, rooted on fish. Bootstrap support values are displayed.

**Dataset S4 (separate file).** Table with results from conservation, acceleration and selection analyses with phyloP and CODEML.
