## Supplementary figures and images for "Broad Host Range of SARS-CoV-2 Predicted by Comparative and Structural Analysis of ACE2 in Vertebrates"

### Fig S3

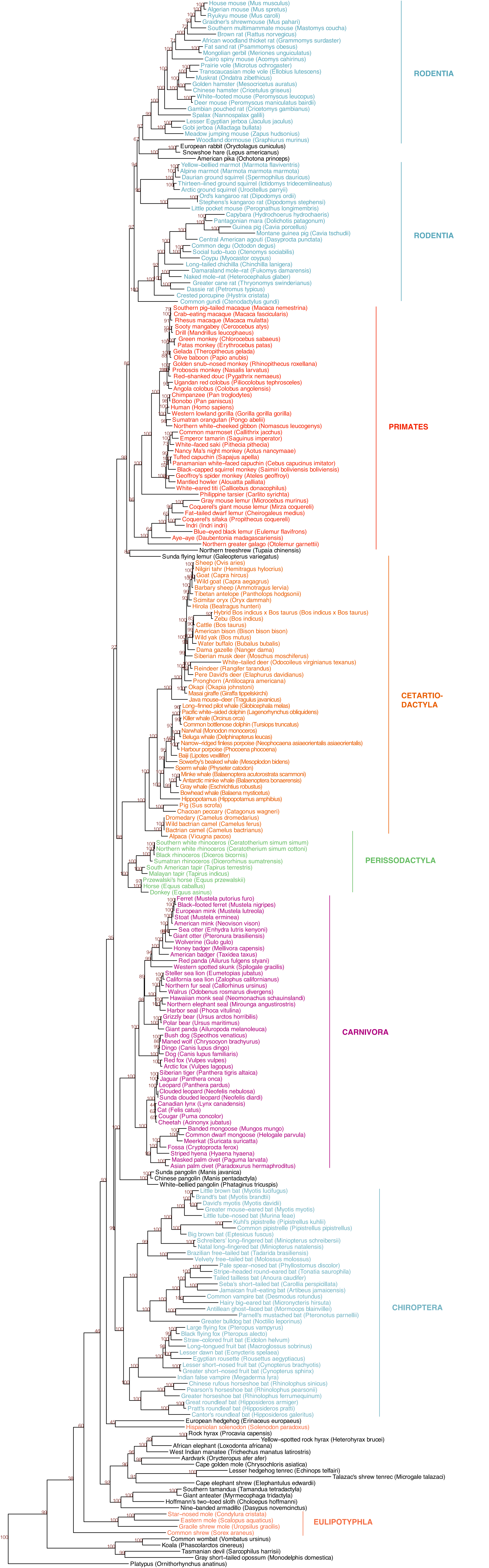
